# Fake it to break it: mimicking superinfection exclusion disrupts alphavirus infection and transmission in the yellow fever mosquito *Aedes aegypti*

**DOI:** 10.1101/2022.09.20.508686

**Authors:** Christine M Reitmayer, Emily Levitt, Sanjay Basu, Barry Atkinson, Rennos Fragkoudis, Andres Merits, Sarah Lumley, Will Larner, Adriana V Diaz, Sara Rooney, Callum J E Thomas, Katharina von Wyschetzki, Kai Rausalu, Luke Alphey

**Author notes:** corresponding author: Luke Alphey.

## Abstract

Multiple viruses cause a phenomenon termed superinfection exclusion whereby a currently infected cell is resistant to secondary infection by the same or a closely related virus. In alphaviruses, this process is thought to be mediated, at least in part, by the viral protease (nsP2) which is responsible for processing the non-structural polyproteins (P123 and P1234) into individual proteins (nsP1-nsP4), forming the viral replication complex. Taking a synthetic-biology approach, we mimicked this naturally occurring phenomenon by generating a superinfection exclusion-like state in *Aedes aegypti* mosquitoes, rendering them refractory to alphavirus infection. By artificially expressing Sindbis virus (SINV) and chikungunya virus (CHIKV) nsP2 in mosquito cells and transgenic mosquitoes, we demonstrated a reduction in both SINV and CHIKV viral replication rates in cells following viral infection as well as reduced infection prevalence, viral titres and transmission potential in mosquitoes.

## Introduction

Superinfection exclusion (SIE) describes a natural phenomenon by which a current infection with one virus prevents a subsequent infection by the same or a closely related virus. It has been described for a range of viruses including human pathogenic viruses, plant viruses and bacteriophages (1–5). One of the groups displaying SIE are alphaviruses (1, 6), a group containing several pathogenic members. Alphaviruses are enveloped, positive-sense RNA viruses belonging to the *Togaviridae* family (7, 8). While some alphaviruses are known to only infect invertebrates (9–11), most members also infect vertebrate hosts and are transmitted between them via arthropod vectors (12). Currently, the most important alphavirus with respect to human health is chikungunya virus (CHIKV) – responsible for multiple disease outbreaks in Africa, Southeast Asia and Americas in recent decades (13).

The exact mechanisms by which certain viruses achieve SIE are either unknown or poorly understood, but there appears to be a wide range of means by which this phenomenon is mediated (4, 5, 14, 15). For alphaviruses, there is evidence that the phenomenon is, at least in part, mediated by the viral protease nsP2 of the primary infecting virus (14, 16). In a natural alphavirus infection, nsP2 processes the non-structural polyproteins P123 or P1234 – translated from the viral genome – in a timely, organised fashion to form the viral replication complex (RC) via intermediate steps crucial to different stages of the alphavirus replication cycle. More specifically, nsP2 mediates cleavage, firstly of nsP3/nsP4, then nsP1/nsP2 and lastly of nsP2/nsP3. Thus, normally it is only after the final processing event that nsP2 is present in its free form and available to mediate SIE against incoming viruses (17–20).

Due to its natural properties, the phenomenon of SIE may present a robust route towards generating alphavirus-refractory systems in mosquitoes. Such a system would be particularly relevant in *Aedes aegypti* – the main vector for CHIKV and other pathogenic alphaviruses (21, 22). Currently there are no approved vaccines available to prevent infection of humans with any pathogenic alphaviruses and the only preventative method is vector control. Control via genetic modification of the mosquito vector to achieve a virus-refractory effect, is a novel approach to overcome this problem (23–25). Of particular interest would be a broad-spectrum effect (i.e. simultaneously affecting multiple viruses) which could potentially be achieved by mimicking a naturally broad-spectrum phenomenon, such as SIE. Recent work suggests that such a route may be feasible: using a trans-replication system (26) Cherkashchenko et al. demonstrated that expression of Sindbis virus (SINV) nsP2 and CHIKV nsP2 was capable of reducing viral RC-dependent reporter expression in an *Ae. albopictus* cell line; the ability of SINV and CHIKV nsP2 to disturbed RC processing of several different alphaviruses including Ross River virus and Mayaro virus (delivered as replicase plasmids) was investigated and both, albeit to varying extents, interfered with RC mediated reporter expression (14).

Here we describe our efforts to employ a synthetic biology approach aimed at developing a virus refractory system in *Ae. aegypti* which functions by mimicking the SIE effect displayed during natural alphavirus infection. We achieve this in both, *Ae. aegypti* cells (Aag2) and transgenic *Ae. aegypti* mosquitoes, through expression of a single viral protein, the viral protease nsP2. Through transiently expressing SINV nsP2 and CHIKV nsP2 in Aag2 cells, we observed a reduction in viral replication of SINV and CHIKV using a trans-replication system. Subsequently, we generated transgenic mosquitoes expressing either SINV nsP2 or CHIKV nsP2 and demonstrated a reduced infection prevalence, as well as significantly lower transmission potential of SINV. Finally, we provide evidence that the expressed nsP2 remains in the cytoplasm in Aag2 cells, the relevant site of action to interfere with the formation of viral RC of the incoming virus.

## Results

### SIE in Aag2 cells using two different fluorescently tagged SINV

To assess if SIE can be detected in *Ae. aegypti* derived Aag2 cells, we infected these cells with mCherry tagged SINV (SINV mCherry) at three different multiplicity of infection (MOI) 0.1, 1 and 10 (or mock-infected) and, after 24h, we infected the same cells with ZsGreen tagged SINV (SINV ZsGreen), at three different MOI for each of the previous treatments. After a further 24h of incubation, ZsGreen fluorescence was measured.

As expected, higher MOIs of SINV ZsGreen resulted in higher levels of ZsGreen expression. At the same time, higher MOI of the primary infecting virus (SINV mCherry) resulted in a greater reduction of ZsGreen fluorescence produced by the secondary infecting virus SINV ZsGreen (Fig 1A). In detail, there was no significant difference (p > 0.05) in ZsGreen expression between cells infected with SINV ZsGreen at MOI 0.1, MOI 1 or MOI 10 alone and cells infected with SINV mCherry at MOI 0.1 prior to the secondary infection with SINV ZsGreen. In contrast, for cells infected with SINV ZsGreen at MOI 0.1, ZsGreen expression is significantly reduced in cells pre-infected with SINV mCherry at MOI 1 (t(10) = 2.998, p = 0.0134) or MOI 10 (t(10) = 3.231, p = 0.0090). Similarly, for cells infected with SINV ZsGreen at MOI 1, ZsGreen expression was significantly reduced in cells pre-infected with SINV mCherry at MOI 1 (t(10) = 5.756, p = 0.0002) or MOI 10 (t(10) = 6.204, p = 0.0001). SINV ZsGreen infection at MOI 10 also resulted in significantly reduced ZsGreen expression in cells pre-infected with SINV mCherry at MOI 1 (t(10) = 5.7565.520, p = 0.0003) or MOI 10 (t(10) = 8.928, p < 0.0001).

**Figure 1:**
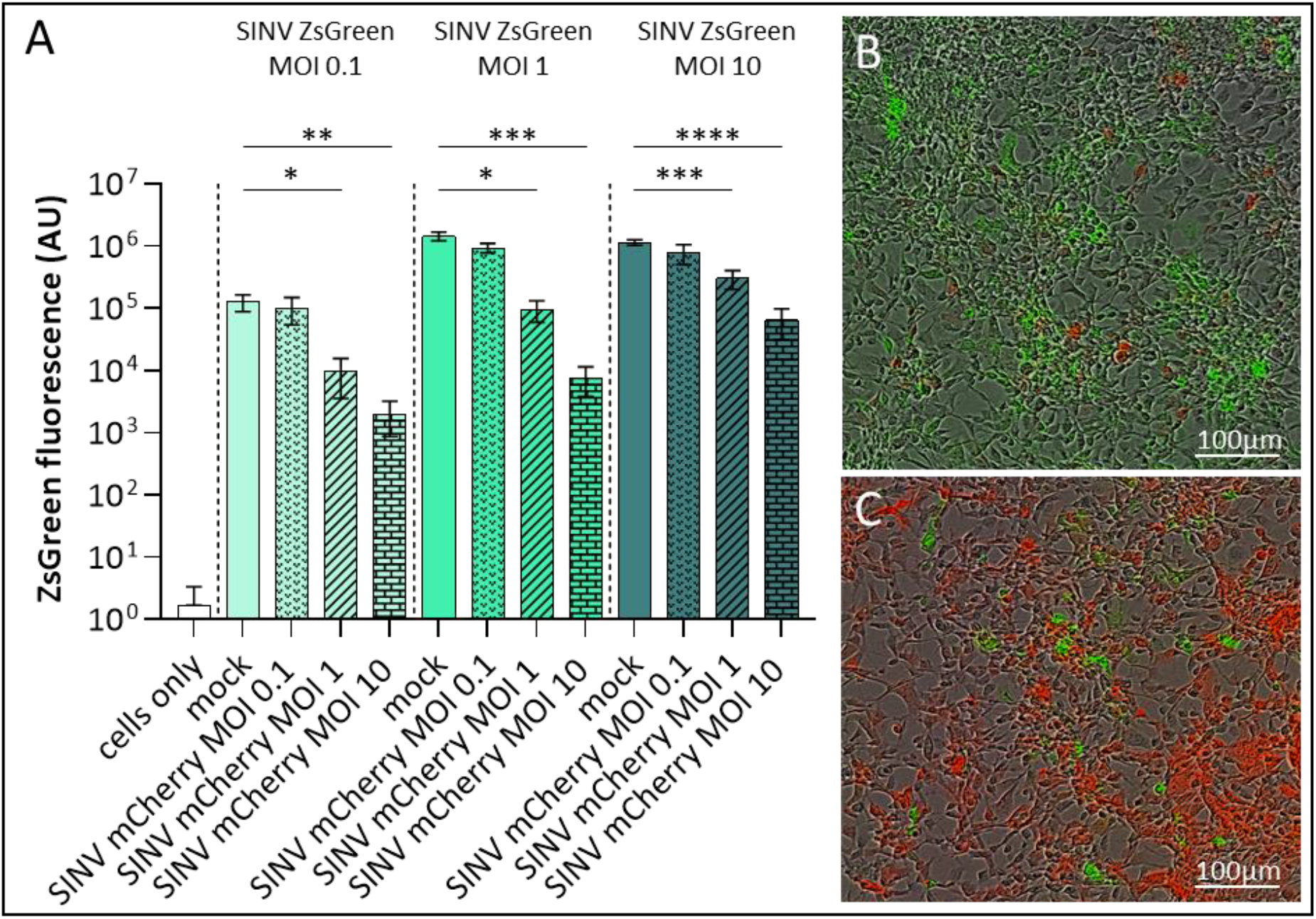
SINV provides strong SIE in Aag2 cells. Aag2 cells were first infected with SINV mCherry at MOIs 0.1, 1 and 10 and control cells were mock infected; 24h later cells were infected with SINV ZsGreen at MOIs 0.1, 1 and 10. (A) ZsGreen expression was measured 24h after infection with SINV ZsGreen using the incucyte cell imaging system as a proxy of SINV ZsGreen viral replication. ZsGreen expression in the absence of SINV mCherry (mock) was measured and, within each SINV ZsGreen MOI group, compared against ZsGreen expression in cells previously infected with SINV mCherry. Significant differences in ZsGreen expression values are indicated with asterisks (*: p<0.05, **: p<0.01, ***: p<0.001, ****: p<0.0001). Representative images showing ZsGreen and mCherry expression in cells infected with SINVmCherry at MOI 0.1/SINVZsGreen MOI 10 (B) and SINVmCherry at MOI 10/SINVZsGreen MOI 0.1 (C).

Images of the cells taken at the same time point show that cells appear to be predominantly infected with either SINV mCherry (red) or SINV ZsGreen (green) (Fig 1B: SINV mCherry MOI 0.1/SINV ZsGreen MOI 10 and 1C: SINV mCherry MOI 10/SINV ZsGreen MOI 0.1). Double-infected cells in which both viruses were replicating express both fluorophores and therefore appear yellow. Yellow appearing cells could be occasionally detected (none shown in Fig 1B or C), thus indicating co-infection of the same cell is possible but presumably rare.

### *In-vitro* reduction of viral RNA replication via nsP2 expression in a split reporter assay

To assess if expression of nsP2 alone can cause a SIE-like state, we designed nsP2 expression plasmids to transiently express nsP2 in Aag2 cells. We tested three different versions of the protease: 1) CHIKV nsP2, 2) SINV nsP2 with a mutation to increase protease activity (nsP2^ND^) and 3) SINV nsP2 with a mutation to abolish the protease activity (nsP2^CA^). In the split reporter activation assay, viral RNA replication activity was measured indirectly, by analysing expression levels of nanoluciferase (nLuc) encoded on an alphavirus reporter template and translated from replicase-generated subgenomic RNA (Fig2A, for details see Methods) (26). Cells were co-transfected with either a SINV or a CHIKV reporter template plasmid, as appropriate for the subsequent virus infection, and CHIKV nsP2, SINV nsP2^ND^, SINV nsP2CA, SINV nsP4 or CHIKV nsP4 expression plasmids. We then measured nLuc levels as a measure of replication activity of SINV or CHIKV in the co-transfected and infected cells and compared to nLuc activity in SINV or CHIKV infected cells only transfected with the respective reporter template.

**Figure 2:**
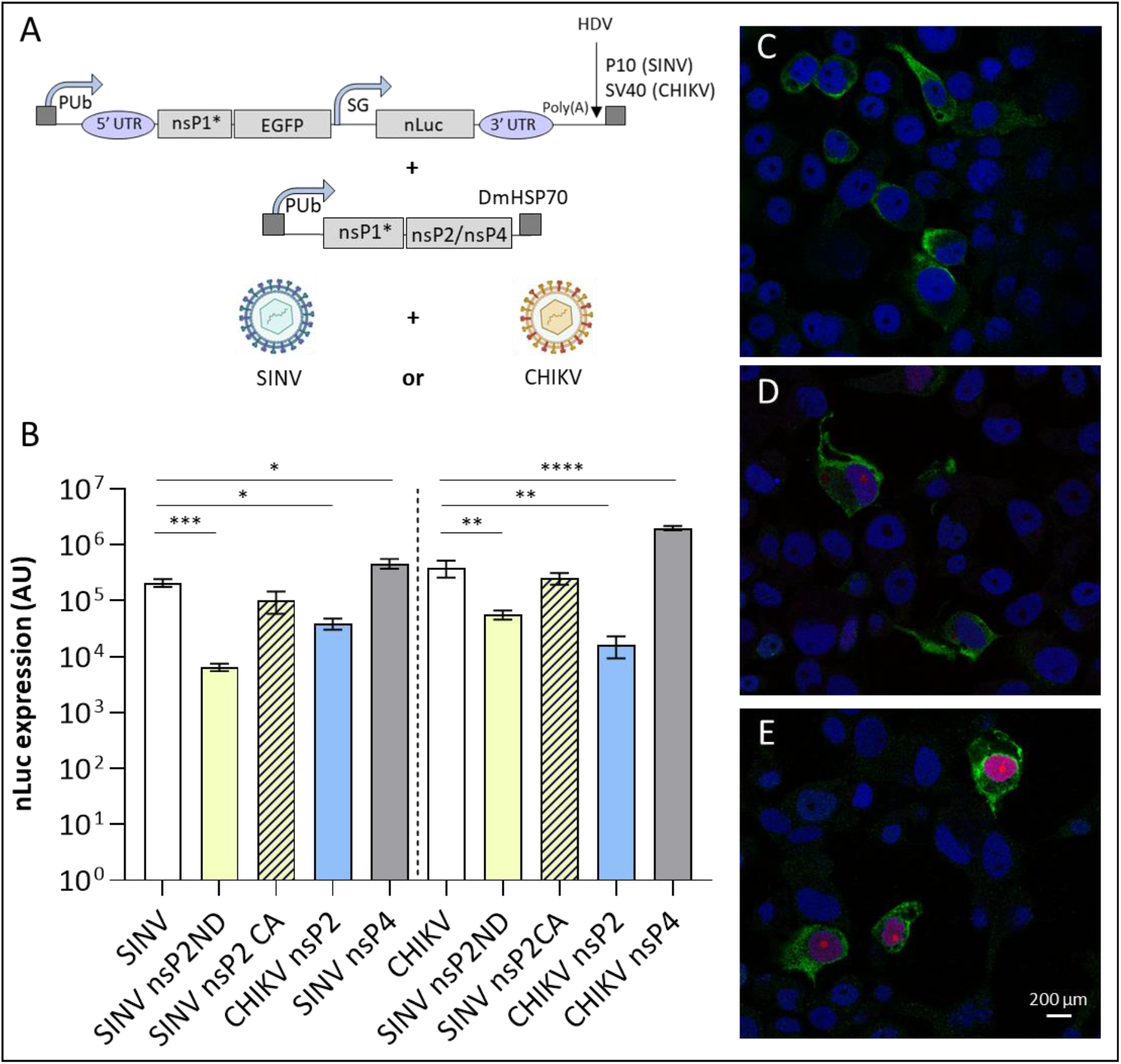
Expression of nsP2 mimics SIE effect in Aag2 cells. (A) Aag-2 cells were co-transfected with a nLuc expressing reporter template plasmid and plasmids expressing different versions of nsP2 or nsP4. (B) 24h post transfection, cells were infected with SINV (left) or CHIKV (right) at MOI 1. Co-expression of fully functional nsP2 (SINV nsP2^ND^ (yellow bars) or CHIKV nsP2 (blue bars) reduced viral replication dependent nLuc expression (nLuc shown as luminescence AU). Co-expression of SINV nsP2 containing a mutation abolishing its protease function (SINV nsP2^CA^ (patterned yellow bars)) led to no reduction in nLuc expression while co-expression of nsP4 (grey bars) cause an increase of nLuc expression compared to no nsP2 control nLuc expression (white bars). Significant differences are indicated with asterisks (*: p<0.05, **: p<0.01, ***: p<0.001, ****: p<0.0001). (C)-(E) Localisation of SINV nsP2 protein in Aag2 cells 24h (C), 48h (D) and 72h (E) after transfection with a SINV nsP2 FLAG expressing plasmid. Localisation of nsP2 in shown in green, DsRed transfection marker in red and DAPI nuclear staining in blue.

Transient expression of functional nsP2 (CHIKV nsP2 or SINV nsP2^ND^) led to a significant reduction in nLuc expression after infection with either SINV or CHIKV compared to expression of the reporter template alone. Transient expression of SINV nsP2CA did not significantly influence nLuc expression while expression of either SINV nsP4 or CHIKV nsP4 led to an increase in expression of nLuc after infection with the respective virus (Fig 2B, left: SINV, right: CHIKV). In detail, co-transfection with SINV nsP2CA did not cause a change in nLuc expression compared to nLuc expression in the absence of nsP2CA expression plasmid neither after infection with SINV nor CHIKV (p > 0.05) in the respective trans-replication system. Both co-transfection with SINV nsP4 and CHIKV nsP4 in the respective trans-replication system did not lead to a reduction in nLuc expression. In fact, co-transfection with SINV nsP4 caused a modest nLuc expression increase (t(13) = 2.464, p = 0.0284); similarly, co-expression of CHIKV nsP4 also caused an increase in nLuc expression (t(9) = 7.391, p < 0.0001). After infection with SINV, there was significantly less nLuc induction in cells co-transfected with one of the two fully functional nsP2 variants; more so with SINV nsP2^ND^ (t(10) = 4.939, p = 0.0006) than CHIKV nsP2 (t(10) = 4.087, p = 0.0022). After infection with CHIKV, similarly to what was observed in the SINV system, there was significantly less nLuc expression in cells co-transfected with one of the two fully functional nsP2 variants. Again, nLuc was reduced to a greater extent homotypically, in this case more by CHIKV nsP2 (t(10) = 3.434, p = 0.0064) than SINV nsP2^ND^ (t(11) = 3.284, p = 0.0073).

### nsP2 is predominantly cytoplasmic in Aag2 cells

To determine the sub-cellular localisation of nsP2 protein in Aag2 cells, we designed a further nsP2 variant, SINV nsP2 FLAG, containing a DYKDDDDK-tag sequence added to the C-terminus of the protein via a flexible linker (SGGSGG). We monitored intracellular localisation of transiently expressed nsP2 in Aag2 cells. At three timepoints (24h, 48h and 72h post transfection) nsP2 was observed localised in the cytoplasm (shown in green, Fig 2C-E). The transformation marker DsRed containing a nuclear localisation signal (NLS) can be seen (Fig 2C-E, in red) only from 48h post transfection due to slow DsRed tetramer formation and maturation (27)) and, as expected, shows a clear co-localization with the nuclear dye DAPI (shown in blue).

### Virus infections of nsP2 expressing transgenic mosquitoes

Transgenic female *Ae. aegypti* expressing either SINV nsP2^ND^ or CHIKV nsP2 driven by an *Ae. aegypti* polyubiquitin promoter (PUb, (28)), were offered an infectious blood meal containing either SINV (1 x10^9^ PFU/ml) or CHIKV (5 x10^8^ PFU/ml), 5-7 days after emergence as adults. At 7 days post infection (dpi), saliva and body samples (not including wings and legs) were collected from each of the mosquitoes. Saliva samples were analysed for presence of virus (transmission potential prevalence), body samples were analysed for presence of virus (infection prevalence) and viral titres of positive body samples were calculated (TCID_50_). Only saliva samples from corresponding virus-positive body samples were analysed for virus prevalence.

After infection with SINV, *Ae. aegypti* Liverpool strain (LVP) females had a significantly higher infection prevalence compared to CHIKV nsP2 expressing females (97% vs. 70%, p = 0.0122) and SINV nsP2^ND^ expressing females (97% vs. 43%, p < 0.0001) (Fig 3A). We did not observe a statistically significant difference in infection prevalence between SINV nsP2^ND^ and CHIKV nsP2 expressing females (43% vs 70%, p = 0.0673). Viral TCID_50_ titres of infected females were determined and a one-way ANOVA revealed that there was a statistically significant difference in SINV titres between the groups (F(2,60) = 15.50, p < 0.0001, Fig 3C). Tukey’s multiple comparisons test found that the mean TCID_50_ values of virus positive LVP females were significantly higher than in CHIKV nsP2 expressing females (p = 0.0053, 95% C.I. = 19332636, 128986320), and SINV nsP2^ND^ expressing females (p < 0.0001, 95% C.I. = 79342130, 207074461) as well as between SINV nsP2^ND^ and CHIKV nsP2 expressing females (p = 0.0440, 95% C.I. = −136575437, −1522198). LVP females had a significantly higher transmission potential prevalence (SINV positive saliva samples as percentage of infected mosquitoes) compared to SINV nsP2^ND^ expressing females (96% vs. 38%, p < 0.0001) but not compared to CHIKV nsP2 expressing females (96% vs. 81%, p = 0.1476, Fig 3E). SINV nsP2^ND^ expressing females also had a significantly lower transmission potential prevalence compared to CHIKV nsP2 expressing mosquitoes (38% vs. 81%, p = 0.0248).

**Figure 3:**
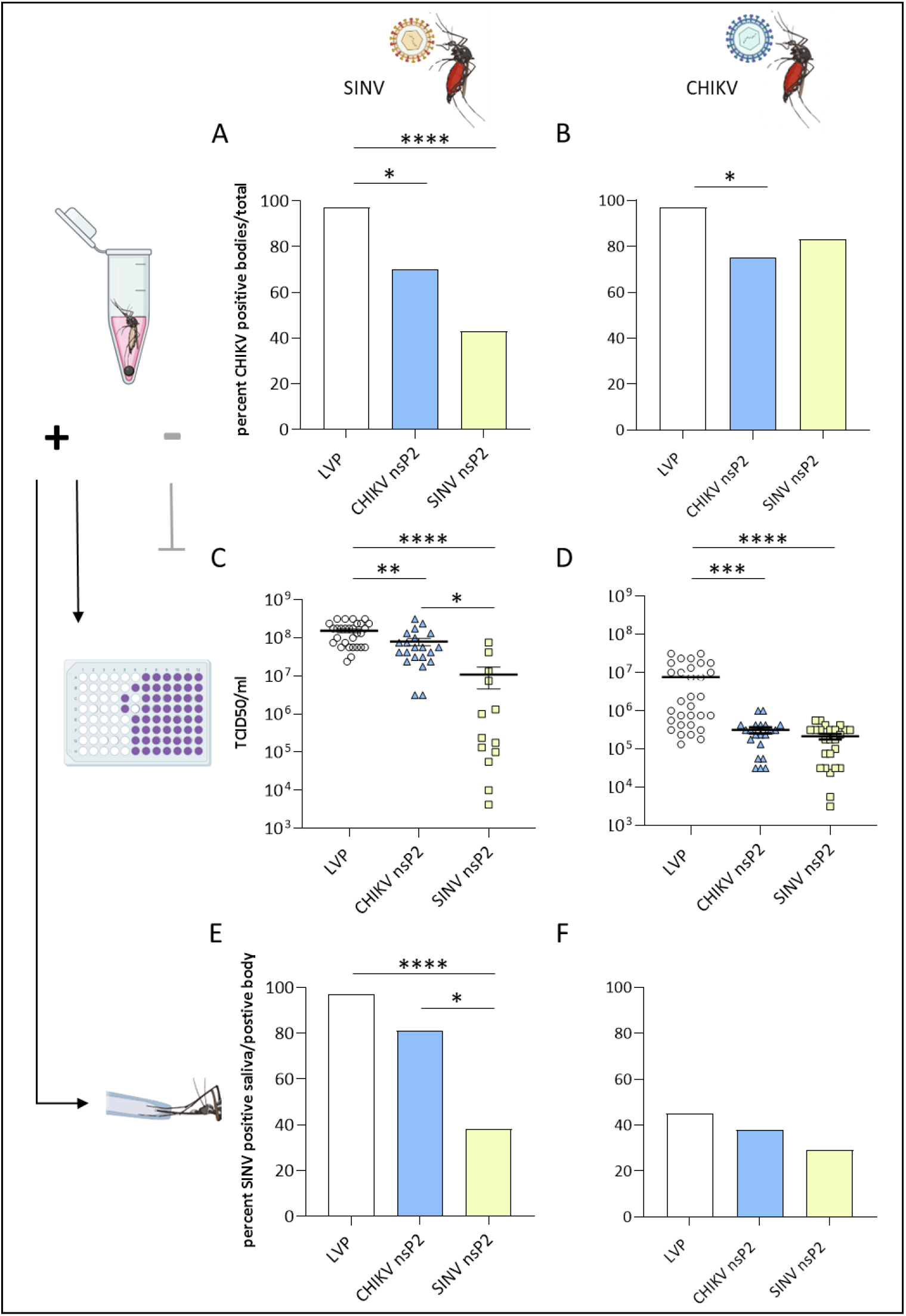
Expression of nsP2 reduces SINV infection prevalence, viral titre and transmission potential prevalence. (A, B) Percent of SINV (A) or CHIKV (B) infected mosquitoes from three different treatment groups are shown, LVP control, CHIKV nsP2 and SINV nsP2 expressing transgenic lines over total number of mosquitoes being provided an infectious blood meal (C, D). SINV (C) and CHIKV (D) TCID_50_ viral titres are shown for the three different treatment groups for virus positive samples. (E, F) Corresponding saliva samples were analysed of positive whole-body samples and are shown as percent over total number of mosquitoes being provided an infectious blood meal. Significant differences are indicated with asterisks (*: p<0.05, **: p<0.01, ***: p<0.001, ****: p<0.0001).

After infection with CHIKV, LVP females had a significantly higher infection prevalence compared to CHIKV nsP2 expressing females (97% vs 75%, p = 0.0201, Fig 3B). We did not observe a statistically significant difference in infection prevalence on day 7 post infection with CHIKV between SINV nsP2^ND^ and LVP or between SINV nsP2^ND^ and CHIKV nsP2 expressing females (p > 0.05). Viral TCID_50_ titres of CHIKV infected females were determined and a one-way ANOVA revealed that there was a statistically significant difference in CHIKV titres between the groups (F(2,74) = 13.01, p < 0.0001, Fig 3D). Tukey’s multiple comparisons test found that mean TCID_50_ values were significantly different between LVP and CHIKV nsP2 expressing females (p = 0.0002, 95% C.I. = 3118995, 11540710) and LVP and SINV nsP2^ND^ expressing females (p < 0.0001, 95% C.I. = 3424895, 11434895) but not between SINV nsP2^ND^ and CHIKV nsP2 expressing females (p = 0.9984, 95% C.I. = −4510246 to 4310161). After CHIKV infection, LVP females appeared to have higher transmission potential prevalence compared to SINV nsP2^ND^ and CHIKV nsP2 expressing females, however, not statistically different (p > 0.05, Fig 3F). There also was no statistically significant difference in transmission potential prevalence between CHIKV nsP2 and SINV nsP2^ND^ expressing female mosquitoes (p > 0.05).

## Discussion

Although SIE is still a poorly understood phenomenon, it has been suggested that in alphaviruses the proteolytic activity of nsP2 of a primary infecting virus plays a part in prohibiting infection of a second virus (6, 29). We have shown that expression of this multifunctional protein reduces replication of two different alphaviruses, SINV and CHIKV, in Aag2 cells. We generated transgenic *Ae. aegypti* lines which express either SINV nsP2 or CHIKV nsP2 and demonstrated that by mimicking a persistent SIE-like state in this mosquito vector, subsequently infected mosquitoes exhibit a reduced infection prevalence, viral titres in infected mosquitoes and transmission potential prevalence, particularly after infection with SINV.

Initially, using an *Ae. aegypti* derived Aag2 cell culture system and two different SINV reporter viruses, we demonstrated that infection with one alphavirus inhibits replication of a second SINV reporter virus in the same cell. Alphavirus SIE had previously been demonstrated in mammalian (30) and other mosquito cell lines such as *Ae. albopictus* C6/36, U4.4 and C7-10 (6). Using fluorescent reporter viruses, we demonstrated that infection with SINV substantially inhibits replication of a second alphavirus (in this case, SINV ZsGreen), consistent with previous work in other mosquito cell systems (6, 31). As here, previous studies of SIE in cell culture did not show a complete prevention of viral replication of the secondary infecting virus (6). However, as analysis in previous studies was carried out by observation of cytopathic effect, it is unclear if replication of the two different viruses occurred in the same, or in different cells. To gain more information, we utilised fluorescent marked viruses and live-cell imaging and found the majority of cells expressed either the reporter of one virus (mCherry - red) or the other virus (ZsGreen - green), rather than both fluorophores (Fig 1B, 1C)). This was indicative of replication of predominantly either, but not both, viruses within the same cell.

After demonstrating alphavirus SIE in Aag2 cells, we investigated whether expression of one viral protein alone, nsP2, could recreate this effect. Previous work in another mosquito cell line had established that expressed nsP2 plays a role in disrupting replicase-delivered RC formation (14). Using a modified trans-replication system (Fig. 2A), we demonstrated that SINV and CHIKV derived nsP2 greatly reduced nLuc expression (henceforth a proxy for viral RNA replication) after infection with either SINV or CHIKV, in both cases to a greater extent when in a homotypic infection scenario (e.g. SINV nsP2/SINV > SINV nsP2/CHIKV). Concurring with this proposed proteolytic SIE mode-of-action, co-transfection with a protease-dead SINV nsP2^CA^ mutant did not significantly affect viral replication. Additionally, co-transfection with either SINV or CHIKV derived nsP4, included as a control to show that inhibition is a specific property of nsP2, actually increased viral RNA replication. This may be attributable to an increased availability of the usually limited nsP4 (due to a naturally occurring opal stop codon at the 3’ end of nsP3 limiting synthesis (reviewed in (32)) combined with rapid degradation of nsP4 protein following the N-end rule (33)) for replication of the used template RNA. Our results are in line with previous work (6, 29) and concur with results obtained by Cherkashchenko et al. (14). However, while there the authors co-transfected plasmids expressing nsP2 and a ns-polyprotein as a source of viral RC, we instead first transfected with nsP2 expressing plasmid and subsequently (24h later) infected those same cells with active virus. This was to provide a more realistic timeline of nsP2 expression as envisaged for virus infection experiments in transgenic mosquitoes. Together with lower plasmid transfection concentrations used in our study (32-60 times lower), the above difference in experimental design might account for variance in effect sizes observed in some of the tested experimental conditions compared to Cherkashchenko et al.

Interestingly, our observations and those of Cherkashchenko et al. (14) differ markedly from another recent study, by Boussier et al., in which no nsP2-mediated SIE effect against CHIKV was observed (8). This difference may be due to nsP2 protease activity depending on the correct N-terminal specification. Here, we ensured faithful recreation of this correct specification by including the DNA sequence encoding the final 10 amino acids of nsP1 upstream of the nsP2 sequence, from which nsP2 can then process itself, mimicking the natural production of nsP2 from a larger polyprotein. Contrastingly, Boussier et al. expressed nsP2 without this nsP1-derived sequence using an artificial start codon. Previous work has shown that an incorrect N-terminal residue or presence of even one additional residue can lead to severely reduced proteolytic activity of nsP2 (34, 35). In addition, Boussier et al. used a mammalian cell culture system and had difficulties generating an nsP2 expressing stable cell line. We did not experience such difficulties in our insect cell culture system when transiently expressing nsP2 and observed a robust SIE-like effect of nsP2 against CHIKV in Aag2 cells. These earlier findings in combination with ours could indicate that SIE against CHIKV might be mediated via different mechanisms in mammalian versus insect systems.

An ongoing question in alphavirus replication, concerns the sub-cellular localisation of free nsP2 and what impact this has on viral replication dynamics. In a mosquito cell line, it was observed that mutations altering the putative NLS of nsP2 did not impact the ability of this protein to reduce RC-dependent replication (14). Our nsP2 confocal imaging data provides an explanation for this observation: we could not detect trans-localisation of nsP2 into the nucleus in Aag2 mosquito cells, indicating that, unlike in mammalian cells, these NLS are non-functional in mosquitoes. This could also potentially explain the absence of a cytopathic effect of nsP2 expression in mosquito compared to mammalian cells – an effect which has complicated functional studies of this important viral protein in mammalian systems (8, 30, 36).

In summary, as a proof-of-principle, we have successfully generated transgenic mosquitoes expressing either SINV nsP2 or CHIKV nsP2. No major adverse fitness effects, e.g. lethality or sterility, were observed in these transgenic mosquitoes, though more detailed studies would be needed to quantitatively assess more subtle effects. Upon viral challenge via infectious blood meal, we demonstrated a significant reduction in infection prevalence, viral titres of those mosquitoes that were infected and prevalence of transmission potential after infection with SINV compared to LVP females. The same trend, albeit not statistically significant for all parameters, was observed upon challenge with CHIKV. Interestingly, results from our cell culture experiments failed to accurately predict a SIE effect of nsP2 in transgenic mosquitoes. While, similarly to cell culture data, we saw a better homotypic than heterotypic SIE effect against both SINV and CHIKV, the SIE effect with respect to infection prevalence against CHIKV in transgenic mosquitoes was weaker than cell culture data would have suggested. In mosquitoes infected with CHIKV, viral titres were greatly reduced, however, not below viral detection threshold. This could be due to a number of different factors, including differences in the evolution of vector/virus interactions – *Ae. aegypti* is one of the main vectors for CHIKV but not SINV, which mainly is transmitted by ornithophilic mosquito species. The observed effect could conceivably be the result of interactions between transgene insertion site-specific tissue expression patterns (we attempted but did not manage to locate insertion sites for either of the two nsP2 expressing transgenic lines) and infection and replication patterns of the respective viruses (e.g. salivary gland lobe specific expression differences between SINV and CHIKV (37–39)).

To our knowledge, this is the first reported study of engineered mosquitoes expressing a viral protein to achieve a viral refractory effect by mimicking a natural viral replication limiting phenomenon, in this case SIE. Alphavirus nsP2 is a highly conserved protein, and its function is crucial to alphavirus replication, making it an ideal target in such refractory systems for avoiding resistance evolution. Furthermore, this conservation may allow broad-spectrum application of identified components. Further work is required to identify key elements of the multifunctional alphavirus nsP2 to improve the identified refractory effect without risking inducing fitness costs to the mosquito. This study, together with recent work (14), provides evidence that an engineered SIE system holds the potential for a broad spectrum, virus-strain independent, low-resistance, synthetic biology approach to develop alphavirus refractory mosquitoes.

## Online Methods

### Construction of nsP2 plasmids

Construction of nsP2 expression plasmids was performed as described in Cherkashchenko et al. (14). In short, sequences encoding full length nsP2 of SINV (isolate Toto1101, (40)) or CHIKV (isolate LR2006OPY1, East/Central/South African genotype) were codon optimized for usage of *Aedes aegypti* and cryptic splice sites present in these sequences were removed. SINV nsP2^ND^contains a single amino acid substitution at position 614 changing aspartic acid (D) to asparagine (N), resulting in an increased protease activity (41). SINV nsP2^CA^ contains a single amino acid substitution at position 481 changing cysteine (C) to alanine (A), resulting in inactivation of the protease activity of nsP2. Expression of the transgene is driven by *Ae. aegypti* polyubiquitin promoter (PUb) (28) and the plasmid contains a transcription terminator of hsp70 gene from *Drosophila melanogaster*, as well as a HR5/iE1 promoter driving expression of the red fluorescent protein DsRed functioning as transformation/transgenic marker. Sequences of all plasmids were confirmed using Sanger sequencing. The correct nsP2 N-terminal residue (glycine for CHIKV nsP2 and alanine for SINV nsP2) is important for its various functions, including its protease activity (34, 35, 42). We therefore encoded not only full nsP2 but also a 10 C-terminal amino acid residue of the respective nsP1 which after translation is removed by nsP2 protease cleavage leaving fully functional nsP2.

### Cell lines

*Aedes aegypti* derived Aag2 cells were maintained in Leibovitz’s L-15 medium (Life Technologies) supplemented with 10% heat-inactivated fetal bovine serum (FBS, Gibco), 10% tryptose phosphate broth (TPB, Gibco), 100 U/ml penicillin (Gibco) and 0.1 mg/ml streptomycin (Gibco) at 28°C. BHK-21 baby hamster kidney cells (CLL-10, ATCC) were cultured in Glasgow’s minimal essential medium (Life Technologies), supplemented with 10% heat-inactivated FBS and 10% TPB and 100 U/ml penicillin and 0.1 mg/ml streptomycin at 37°C and 5% CO_2_.

### Virus stocks

SINV AR339 stocks were propagated in BHK-21 cells by culturing for 48h at 37°C. SINV (Toto1101) harbouring either a mCherry or ZsGreen marker and a duplicated subgenomic promoter in the intergenic region and CHIKV (isolate LR2006OPY1) were rescued from icDNA clones. icDNA plasmids containing SP6 promoters, were linearized, in-vitro transcribed using the mMESSAGE mMACHINE™ SP6 Transcription Kit (Thermo Fisher) and the resulting capped RNA electroporated into BHK-21 cells using a Gene Pulser Xcell electroporator (BioRad) and cultured for 48h at 37 °C. The culture media was harvested, clarified by centrifugation. Virus stocks used for mosquito infection studies were concentrated using Amicon™ 100 kDa Ultra-15 Centrifugal Filter Units (Merck) to achieve appropriate virus concentrations in the resulting blood meal. Virus stocks were aliquoted and stored at −80 °C.

### SIE exclusion in Aag2 cells

To test whether SIE can be detected in *Ae. aegypti* cells, Aag2 cells were infected with SINV mCherry at three different MOI (0.1, 1, 01) and after 24h of incubation a secondary infection with SINV ZsGreen was carried out. For each of the infections, the inoculum was left on the cell monolayer for 1h and subsequentially exchanged for fresh L-15 media. Plates were imaged using the IncuCyte live cell imager (Sartorius) after 24h of incubation following the secondary infection and images analysed using the associated IncuCyte software (Sartorius). The experiment was carried out in eight technical replicates per treatment.

### Trans-replication nsP2 assay

Three different versions of nsP2 protease were tested: 1) CHIKV nsP2, 2) SINV nsP2 with a mutation to increase protease activity (nsP2^ND^) and 3) SINV nsP2 with a mutation to abolishing the protease activity (nsP2^CA^). The N614D mutation was chosen as previous work identified this mutation in SINV to enhance protease activity and disrupt viral replication (41). Additionally, more recent work using a trans-replicase approach (14) found that this mutation in transiently expressed SINV nsP2 delivered a slightly stronger effect compared to wild-type SINV nsP2 in a different mosquito cell line. The C481A mutation of SINV nsP2 was added to the panel to test if the protease function of nsP2 is needed to achieve a SIE effect. SINV/CHIKV nsP4 expression plasmids were added to the panel to function as a negative control accounting for any non-specific effect that the expression of a viral non-structural protein, or a plasmid of this structure, might have on viral replication.

In this split reporter activation assay, viral replication activity was measured indirectly by quantifying expression levels of a nanoluciferase (nLuc) marker encoded on an alphavirus reporter template following the subgenomic promoter (Fig 2A). In this system the reporter template resembles the positive-sense viral genome RNA containing the 5’ and 3’ UTR as well as the subgenomic promoter of a specific alphavirus. Non-structural and structural proteins are exchanged for reporters. The reporter RNA cannot replicate itself but provides a substrate which can be replicated in trans by viral RNA replicase. Upon viral infection of a cell, viral components of the replication complex will be expressed; these will act on the reporter RNA leading to transcription and translation of the subgenomic RNA encoded nLuc reporter protein (26).

As the reporter plasmid and the nsP2 plasmid are co-transfected into the same cells, this system is robust to the low transfection efficiency commonly seen in Aag2 cells (26). Cells were co-transfected with either a SINV or a CHIKV reporter template plasmid (as appropriate for the subsequent virus infection) and CHIKV nsP2, SINV nsP2^ND^, SINV nsP2^CA^, SINV nsP4 or CHIKV nsP4 expression plasmids. We then measured nLuc levels as a measure of replication activity of SINV and CHIKV in the co-transfected cells.

Aag2 cells were seeded into 96-well plates (55,000 cells/well). After 24h, cells were co-transfected with the viral reporter template plasmid appropriate for the virus to be tested (10ng/well), one of the plasmids in the panel to be tested (encoding for SINV nsP2^CA^, SINV nsP2^ND^, CHIKV nsP2, SINV nsP4, each 20ng per well) and a firefly luciferase expressing plasmid (HR5/IE1-firefly-SV40, 0.2ng) as internal control. The used trans-replication assay and reporter template plasmids have been described elsewhere (14, 26, 43). Transcription of both SINV and CHIKV reporter template plasmids is driven by a truncated *Ae. aegypti* PUb promoter and the plasmids contain either a SV40 (CHIKV) or P10 (SINV) transcription terminator (Fig 2A). Transfection was performed using TransIT-PRO® Transfection Reagent & Kit (Mirus Bio). Post transfection, cells were incubated at 28°C and infected with the respective virus inoculum at MOI1 24h later. After a further 48h of incubation cells were harvested and luciferase assays were performed using the Nano-Glo® Dual Reporter Assay Kit (Promega) and the GloMax®-Multi Detection luminometer System (Promega). The experiment was carried out in 8 technical replicates per treatment.

### Immunohistochemistry

Alphaviruses display a cytopathic effect in mammalian cells. This is thought to be mediated, at least in part, by translocation of a fraction of nsP2, once cleaved off of the P23 polyprotein precursor, into the nucleus and there causing host cell transcriptional shut off (30, 36) Translocation of free nsP2 protease is mediated by nuclear localisation signals (NLS) within the protease region (44). Mutations disrupting the nuclear localisation signal have been shown to result in reduced cytotoxicity (14, 45, 46). Mosquito cells do not typically display a cytopathic effect upon infection, similarly, infected mosquitoes do not display overt symptomatic disease upon infection. We investigated whether we could find evidence of translocation of transiently expressed nsP2 into the nucleus. Our hypothesis that nsP2 mediates a SIE-like state by prematurely processing virus encoded P123/P1234 of an infecting virus requires sufficient nsP2 to be present in the cytoplasm, so eliminating nuclear localisation might increase the effect.

Aag2 cells were seeded in 24-well plates (120,000 cells/well), with each well containing a round cover slip. Cells were transfected and incubated as described above with a PUb driven, FLAG-tag SINV nsP2 expressing plasmid. After 24h, 48h and 72h a subset of cells was fixed using 10% formalin solution (Sigma) for 1h. After fixation, cells were washed three times with PBS containing 0.1% Tween (PBST, Sigma) and unspecific binding sites were blocked with 3% bovine serum albumin solution in PBST (Sigma). Cells were incubated in mouse anti-DDDDK (FLAG)-tag antibody solution (1:500, Abcam) overnight at 4°C. After three washes in PBST, cells were incubated in anti-mouse Alexa 488 antibody solution (1:200, Abcam) for 1h at RT, washed again twice and mounted using VectaShield Hard set mounting medium containing DAPI (2bScientific). Imaging was carried out using a STELLARIS 5 confocal microscope (Leica) at The Pirbright Institute Bioimaging facilities.

### Generation of transgenic mosquitoes

Transgenic mosquito lines were generated as previously described (47–49). Briefly, *Ae. aegypti* Liverpool strain (LVP) pre-blastoderm embryos were microinjected with a mixture of plasmid expressing hyperactive piggyBac transposase under control of PUb promoter (300 ng/μL) and donor piggyBac transgene plasmid (SINV nsP2^ND^ and CHIKV nsP2, 500 ng/μL). Injection survivors were backcrossed to LVP individuals and progeny screened for the fluorescent transformation marker DsRed using a fluorescent stereomicroscope (Leica) to identify transformed individuals. Single male transformants (G1) were then crossed to five LVP females each to establish independent transgenic lines. For each of the lines used for data presented in Fig 3, approximately half of the offspring inherited the fluorescent marker, indicating single insertions. Insertion-site identification was performed as previously described (50); however, repeated efforts to determine this sequence were unsuccessful. To generate lines with a higher allele frequency for SINV nsP2 and CHIKV nsP2, transgenic individuals were crossed to transgenic individuals only, for a minimum of five generations. Transgene frequency in the lines used for virus challenge studies was between 85-90%.

Transgenic and LVP *Ae. aegypti* were reared and maintained at 26.5°C (±1°C) and 65% (±10%) relative humidity with a 12:12 hour light:dark cycle. Larvae were fed on finely ground TetraMin Ornamental Fish Flakes (Tetra GmbH), and adults on a 10% sucrose solution. Adult females were fed on defibrinated horse blood (TCS Biosciences).

### SINV and CHIKV infection of mosquitoes and virus determination

All experiments with CHIKV were performed under Containment Level 3 (CL-3) conditions at The Pirbright Institute, UK. CHIKV LR2006OPY1 and SINV AR339 passaged in BHK-21 cells were used. LVP or transgenic (confirmed by red fluorescence body marker) female mosquitoes were allowed to feed on an infectious blood meal containing either SINV (1 x10^9^ PFU/ml) or CHIKV (5 x10^8^ PFU/ml) via a temperature-controlled blood feeding system (Hemotek Ltd.) inside a Microbiological Safety Cabinet class III. Mosquitoes were allowed to feed for 1h, immobilised on ice and engorged females were transferred into maintenance containers and maintained for 7 days at 27°C and 12:12 light/dark cycle. For transmission potential prevalence, mosquitoes were immobilised on ice at 7 dpi and wings and legs removed. Saliva was collected according to published methods (51) and the remaining body was collected. Infection prevalence and infection titres were analysed using TCID_50_ assays, transmission potential prevalence was analysed using cytopathic effect assays, both carried out on BHK-21 monolayers. Infection prevalence was defined as percent of positive samples over all investigated samples, infection titres were defined as TCID_50_ titres of all positive samples and transmission potential prevalence was defined as the percent of positive saliva samples out of the number of positive body samples. Experiments were carried out with 30-35 female mosquitoes per treatment.

### Statistical analysis

Analysis of SIE of SINV ZsGreen by SINV mCherry was carried out using Student’s two-tailed t-tests of combinations of interest. Analysis of SIE caused by transient expression nsP2 on SINV and CHIKV infection in Aag2 cells was carried out using Student’s two-tailed test to compare treatment groups of interest. Analysis in mosquito infection studies was carried out using Fisher’s exact tests for infection prevalence and transmission potential prevalence and one-way ANOVA for infection titres. All analysis was carried out using GraphPad Prism 9.3.1.

## Author Contributions

Methodology: BA, CMR, LA, RF, SB and SL; Validation: BA, CMR, RF, SB and SL; formal analysis: CMR; investigation: CMR and EL; sample preparation and data collection: AVD, BA, CJET, EL, KW, SB, SL, SR and WL; writing (original draft preparation): CMR; writing (review and editing): AM, CMR, LA, and SB; visualization: CMR; project supervision: CMR, LA, RF and SB; resources: AM and KR; funding acquisition: AM, LA and RF. All authors have read and agreed to the submitted version of the manuscript.

## Acknowledgments

The authors thank Jennifer Porter, Miles Nesbit and Owen Vaughan for their help with mosquito maintenance and assistance with attempts to identify transgene insertion sites of transgenic mosquitoes. Confocal imaging was carried out at the Bioimaging facilities at the Pirbright Institute. We are grateful to John Fazakerley and Jamal I-Ching Sam for discussions and advice.

